# Positive and negative frequency-dependent selection acting on polymorphism in a palatable moth

**DOI:** 10.1101/2023.04.13.536688

**Authors:** Riccardo Poloni, Marina Dhennin, Johanna Mappes, Mathieu Joron, Ossi Nokelainen

## Abstract

Camouflage and warning signals are contrasted prey strategies reducing predator attack, which offer an excellent opportunity to study the evolutionary forces acting on prey appearance. Edible prey are often inconspicuous and escape predation by remaining undetected. Predators learn to find the most common ones, leading to apostatic selection (advantage to rare morphs) enhancing variation in cryptic prey. By contrast, defended prey are often conspicuous and escape predation by using warning colorations identifying them as unprofitable. Predators avoid the ones they are most familiar with, leading to positive frequency-dependence and warning signal uniformity. It is less clear, however, what happens when two morphs of the same species vary strongly in conspicuousness, and how to explain the maintenance of cryptic and conspicuous morphs within populations, in the case of profitable prey. Using the white and melanic morphs of the invasive Box Tree Moth (*Cydalima perspectalis*) presented at three different frequencies, we investigate whether a) caterpillars and adult moths are palatable for birds, b) the less conspicuous, melanic morph experiences lower predation rates and b) whether frequency-dependence balances morph frequencies. Our results suggest that the melanic morph enjoys a survival advantage owing to a lower visibility. However, our experiments show that, unexpectedly, the two color morphs experience opposite patterns of frequency-dependent predation, despite being both fully palatable to birds. The melanic morph is under apostatic selection, whereas the conspicuous, white morph is subject to positive frequency-dependence (safety in numbers). Our experiments also show some level of unpalatability in the caterpillars. These results offer novel insight into how predation triggers contrasting evolutionary patterns in a palatable, polymorphic species within two morphs that differ markedly in conspicuousness and within two different life stages.

**Lay summary:** Understanding the factors influencing character variation in natural populations is a key question in evolutionary ecology. Predation is one of the main drivers of color evolution in prey communities and prey usually mitigate predation using camouflage or warning colors. Camouflage evolves because it lowers the probability of being detected by predators. Since predators are more efficient at finding prey which they are familiar with, prey which display a rare phenotype are favoured (negative frequency-dependent selection). By contrast, aposematism is defined by conspicuous appearance in toxic or otherwise unprofitable prey, and evolves because birds identify defended prey by learning to use their appearance as a warning signal. The most common signals are usually best identified and avoided (positive-frequency dependent selection). It is not clear, however, how these two forces combine when predators are facing cryptic and conspicuous morphs of the same species, and how to explain their coexistence. Here we investigate this question in a laboratory experiment, by presenting wild birds with a melanic and a white morph of the same moth. Unexpectedly, our results show that despite being both fully palatable to birds, the two color morphs are subject to very different types of selection depending on their frequencies. The melanic morph is favored when it is rare, the conspicuous white morph as it gets common. The simultaneous action of these forces may contribute to maintain color polymorphism in natural populations. We also show that caterpillars of this species are unpalatable and chemically defended, whereas adults are not, showing opposite strategies of predator defense in different life stages of the same species.

## INTRODUCTION

Animal coloration has been the focus of intense research in evolutionary biology since the times of Darwin and Wallace, and has played a pivotal role in our understanding of the links between ecology and evolution (Poulton, 1890; Majerus, 1998; Cuthill et al., 2017). Camouflage evolves in response to predation, because it lowers the probability for a prey to be detected and attacked due to its lower detectability (Merilaita et al., 2017). Background matching, disruptive coloration, or masquerade (resemblance to an inedible object) are the three main types of camouflage (Endler, 2006; Schaefer and Stobbe, 2006). Polymorphism in camouflage is widespread and some taxa can be extremely variable in their appearance, such as in the underwing moths (*Catocala* spp.) where 40% of the species are polymorphic (Bond, 2007). A classic explanation for polymorphism in cryptic prey rests on predator cognition: predators find cryptic prey more easily when they are familiar with their appearance (“search images”), and so attack the common morphs at higher rates (Bond and Kamil, 2002; Bond, 2007). This may lead to negative frequency dependent (or apostatic) selection that favours morphs as they become less common. This scenario contrasts with positive FDS (frequency-dependent selection), when the probability for a prey to be attacked decreases as it becomes more common. Notably, predators learning to associate distasteful prey with their appearance avoid prey they are familiar with, hence favouring the commonest forms (Lindström et al., 1997; Ruxton et al., 2004). Such positive FDS leads to local monomorphism, and is the basis of mimicry among species (Kapan, 2001; Chouteau et al., 2016). However, although positive FDS on aposematic prey and negative FDS on cryptic prey are widely agreed processes, polymorphism in mimicry is in fact quite widespread and the classic association of palatability, conspicuousness and frequency-dependence has many flavours and exceptions (Kapan, 2001; Rönkä et al., 2020). Here, we investigate how selection may act on two colour morphs of a palatable prey when they differ markedly in their conspicuousness.

The Box Tree Moth, *Cydalima perspectalis* (Crambidae), is an moth native to subtropical Asia, which has invaded the whole Europe and part of Palaearctic Asia and North America since its introduction in 2007 in southern Germany (Bras et al., 2019). It displays a marked wing colour polymorphism, with the coexistence of a conspicuous white morph and a grey-brown melanic morph, maintained at about 5:1 ratio (Fig. 1). The genetic and ecological drivers that underlie the maintenance of this polymorphism are still unknown. Bird predation is reported on both adults and caterpillars (Leuthardt et al. 2013; Brua, 2014), and although caterpillars accumulate alkaloids in their tissue, adults appear to be devoid of chemical defences (Leuthard et al. (2013).

**Fig. 1:**
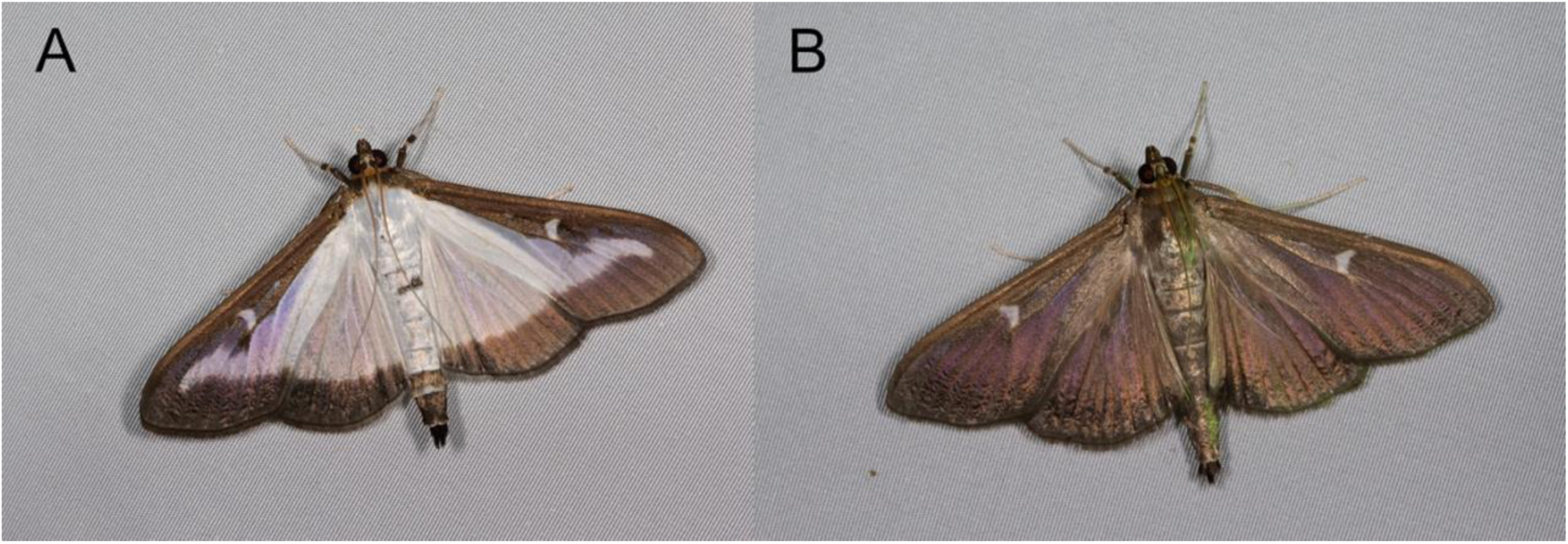
The two different morphs of the Box Tree Moth: white (A) and melanic (B).

The polymorphism in this apparently palatable moth therefore makes it an excellent model to ask whether polymorphism is maintained by negative frequency dependence, or whether the two morphs are subject to distinct selection regimes. Using predation experiments with wild birds maintained in the lab, we first test for palatability variation in adult moths and caterpillars, and then for crypsis and frequency-dependent predation in the two morphs. We show that both morphs are palatable and that the differences in conspicuousness are associated with frequency dependence acting in opposite directions, bringing new insight into the maintenance of prey polymorphism.

## MATERIALS AND METHODS

### Model organisms

Box tree moth specimens were obtained from a laboratory stock founded in 2020 at the Centre d’Ecologie Fonctionelle et Evolutive (France) using wild populations from Saint-Clément-de-Rivière, France (GPS coordinates: 3.84, 43.71) and Le Caylar, France (GPS coordinates: 43.86, 3.32). Wild blue tits (*Cyanistes caeruleus*) were caught from baited traps at Konnevesi Research Station, where predation experiments took place, in February-March 2022. Once trapped, birds were measured, weighted, aged, sexed, and housed individually in plywood cages (80 cm x 65 cm x 50 cm) with food and water *ad libitum*. Wild birds were used with permission from the Central Finland Center for Economic Development, Transport, and Environment, licensed from the National Animal Experiment Board (ESAVI/9114/ 04.10.07/2014) and the Central Finland Regional Environment Center (VARELY/294/2015), and used according to the Association for the Study of Animal Behaviour guidelines for the treatment of animals in behavioural research and teaching.

Details on bird training and maintenance can be found in supplementary methods.

### Palatability tests

Before the experiment, birds were placed in a clean cubicle (80 cm x 65 cm x 50) with water ad libitum and food-deprived for one hour. The experimental design, as described by Winters et al. (2021), allows detecting when the bird sees the proposed items, and therefore measuring the time spent deciding to attack, without any confusion with the time spent searching (Fig. 2A). Each bird was trained with crushed peanuts to get familiar with the experimental system. Once fear and hesitation behaviour were not observed (see further), the bird was presented with 6 randomly chosen prey items to be tested: 3 moths per morph for the tests on adult moths, or 6 randomly chosen caterpillars for the test on larvae. For each prey offered a) the type of prey, b) *hesitation time*, c) *attack time* and d) eating behaviour were recorded. Stress behaviour like beak rubbing was also recorded. *Hesitation time* is defined as the time from the presentation of the prey on the movable tray to the moment when the bird comes onto the tray. A*ttack time* is the time taken by the bird to approach and peck/eat the prey after seeing it, and was used as a proxy for hesitation to attack, following methods developed earlier to evaluate prey palatability e.g. (Exnerová et al., 2015; Rojas et al., 2017). Four categories of eating behaviour were recorded: “eat” (prey eaten completely), “half” (prey partially eaten), “catch” (prey caught but not eaten) and “refuse” (prey not caught).

**Fig. 2:**
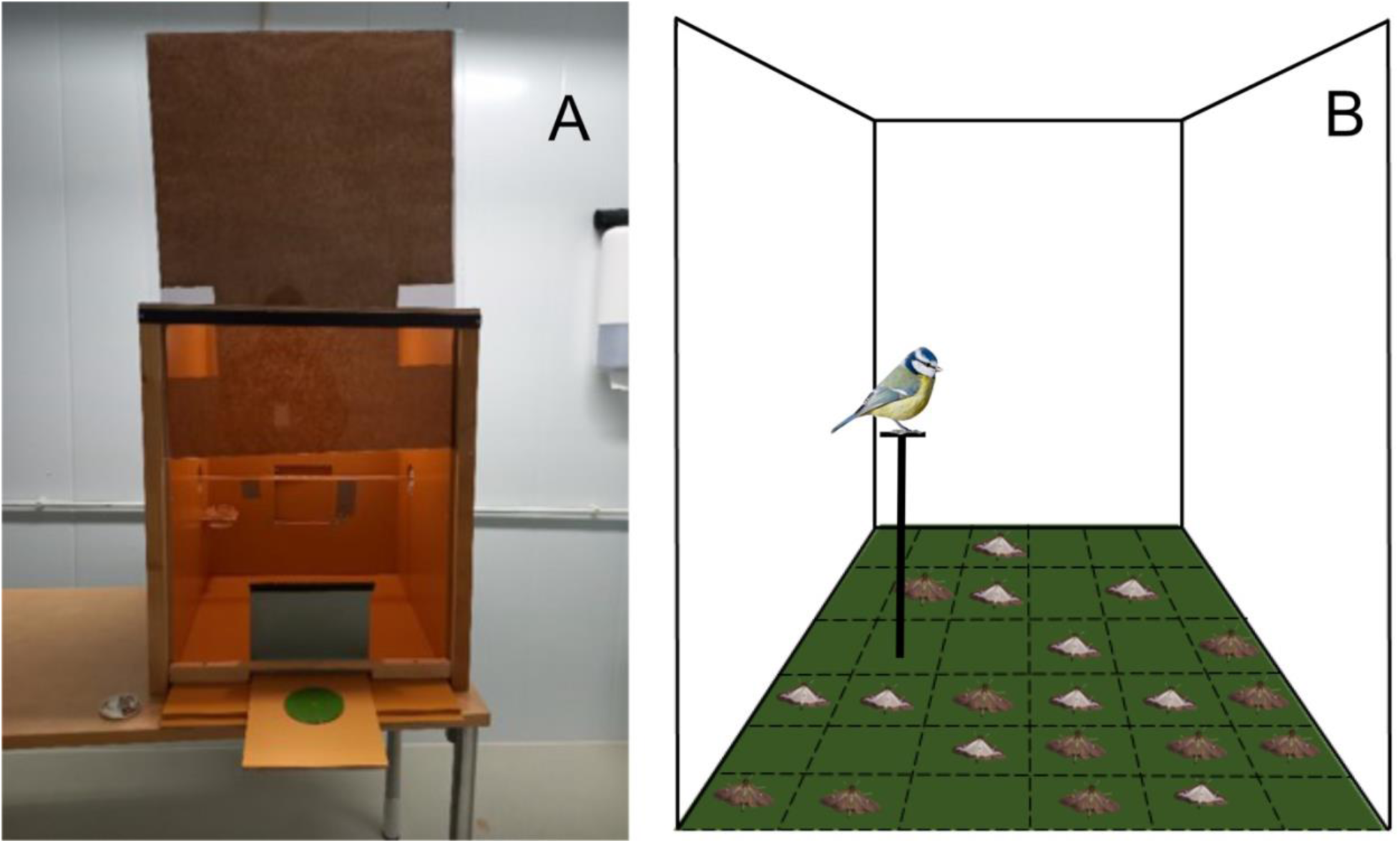
Experimental setup used for palatability assays (A) and predation assays on adult moths (B). Pane A shows the movable tray that allows the bird to see the prey presented on the green plate only when onto the perch.

### Visual modelling

Spectrometer measurements and visual modelling were used to model how birds perceive the background chosen and the two morphs of the Box Tree Moth. Reflectance spectra of white and black wings, camouflage net, and box tree leaves, were obtained using a spectrometer and analysed to detect achromatic and chromatic contrasts. Details on the measurement procedures and analysis can be found in supplementary methods.

### Predation tests

Experiments took place in an aviary (2,80 x 3,70 x 2,20 m^2^, H x L x W) using one bird per trial. The ground was covered with a camouflage net to mimic a heterogeneous, natural background and divided into a grid to record the position of prey items (Fig. 2B). For each trial, 30 moths were placed on the ground with different morph ratios according to three different treatments: control: 1:1 white:melanic; melanic biased: 1:4, and white-biased: 4:1. Each trial ended when birds ate 15 moths.

### Statistical analysis

Statistical analyses were performed using R (3.6.2).

### Palatability

*Attack time* (i.e., the time taken by the bird to approach and attack the prey after seeing them) was used as a proxy for measuring the time the bird spends to attack after seeing the prey. Therefore, we tested for a difference in the *attack time* between control (peanuts) and experimental prey (adult moths or caterpillars). All peanuts offered at the beginning of a trial were removed before analysis. One bird (B36) refused nearly all prey items including peanuts and was removed from all analyses.

**For adults,** *attack time* was tested for normality, using the Shapiro-Wilk normality test. Since the data are not normally distributed, we tested for differences in attack time using a Wilcoxon signed-rank test, with three comparisons: White-Control, Melanic-Control and White-Melanic.

**For caterpillars,** since the refuse rate was high, all the birds that refused all the caterpillars were first removed, and a Wilcoxon signed-rank test was then used to compare attack times of caterpillars and controls. Second, to include the whole behavioral variation and all the birds, a Wilcoxon signed-rank test was used to compare refuse rates of caterpillars and peanuts, and without removing birds that refused all caterpillars. In both cases, data were not normally distributed.

A linear model was implemented to test whether birds with lower weight accepted caterpillars at a higher rate (refuse rate ∼ weight). Finally, generalized linear mixed models were fitted to assess the effect of different variables on the refuse rate, with bird behaviour (eat/refuse) as response variable and logit link. Bird age, sex and trial duration were included as fixed effects. Bird identifier was included as a random factor to account for variance in bird behaviour. Only significant variables (p<0.05) were retained for subsequent models. All models were then compared using Akaike Information Criteria.

### Predation

To measure if different colour morphs were more attacked, we focus on two different measures of predation. The *overall attack rate* was defined as the number of attacked moths of a certain morph over all available moths presented, and the *morph-specific attack rate* as the number of attacked moths of a certain morph over available moths of that same morph. In a random scenario, assuming no difference in detectability between morphs and an absence of frequency-dependent predation birds would be expected to attack the moths according only to their abundance. The observed *morph-specific attack rate* of each morph was then compared with the expectation (0.5) using a one-sample t-test (see also explanation in Fig. 5). Finally, the different treatments were compared using a Welch two-samples t-test to check if proportions of eaten moths changed with different frequencies.

As for the palatability tests, the effect of different variables on attack rate was tested for by fitting generalized linear mixed models with the variable found/not found as response variable under a binomial distribution and logit link. Morph and treatment were included as fixed effects as well as their interactions. Bird age, sex and prey grid position (edge or centre) were also included as fixed effects. Since trials ended when birds had eaten a fixed number of 15 moths, trial duration was added as a covariate. Bird identifier was included as a random factor to account for variance in bird behaviour. First, a model was performed including all variables, then, only grid position and morph emerged as significant variables explaining attack probability and were thus retained, together with treatment, which was still retained. Overall, the model with lowest AIC resulted the GLM including morph, treatment and grid position (edge or centre) as fixed factors (AIC = 1835, Table 1).

**Table 1:** Fixed effects for the generalized linear model performed with predation data, including morph, treatment and grid position (edge or centre) as fixed factors (AIC = 1835).

|  | **E** | **SE** | **Z** |
| --- | --- | --- | --- |
| **(Intercept)** | 0,736 | 0,152 | 4,848 |
| **Butterfly colour (M)** | -0,536 | 0,128 | -4,185 |
| **Treatment (50:50 W:M)** | -0,158 | 0,141 | -1,119 |
| **Treatment (80:20 W:M)** | -0,328 | 0,156 | -2,096 |
| **Position (edge)** | -0,556 | 0,111 | -4,989 |

Data tables with raw data and the scripts used are available in supplementary materials and at https://github.com/r-poloni/predation (after publication).

## RESULTS

### Palatability

Palatability data for adult moths were obtained using 266 individual prey presented to 27 birds tested. Out of 80 white and 78 melanic moths presented, 76 and 77 were eaten (respectively). Average *attack time* was 43 s for peanut, 20 s for the melanic moth and 14 s for the white moth. *Attack time* did not follow a normal distribution (Shapiro-Wilk normality test). No significant difference in *attack time* was observed between moths and peanut (Wilcoxon signed-rank test: control/melanic morph, p = 0.5153, control/white morph, p = 0.1731) or between moth morphs (Wilcoxon signed-rank test: white morph/melanic morph, p = 0.1899) (Fig. 3).

**Fig. 3:**
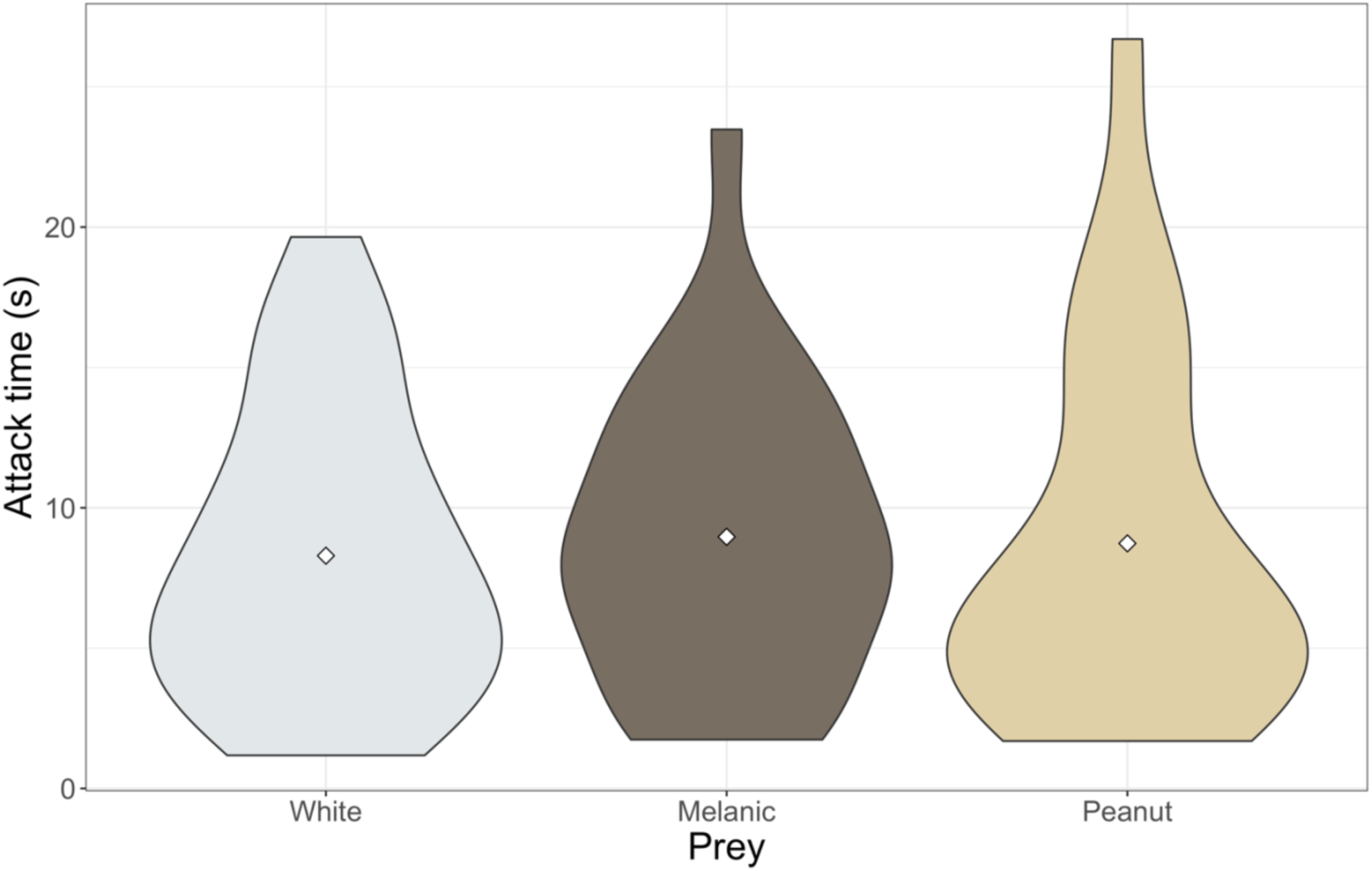
Violin plot showing the attack time in the palatability assay on adult moths. No difference in *attack time* (i.e., the time taken by the bird to approach and peck/eat the prey after seeing) was observed between moths and peanuts (Wilcoxon’s signed-rank test, control/melanic morph p = 0.5153, control/white morph 0.1731) or among moth morphs (white morph/melanic morph, p = 0.1899).

Palatability data for caterpillars were obtained using 458 prey (peanuts + caterpillars) presented to 40 birds. Out of 221 caterpillars presented, 73 were refused, 40 attacked but not eaten, 22 half-eaten, and 86 eaten. Refuse rate and eating behaviour varied greatly among birds (Fig. 4). *Attack time* was markedly higher for caterpillars than for peanuts (Wilcoxon signed-rank test: control prey/caterpillars, p < 0.001.) (Fig. 4). Refuse rate differed markedly between peanuts (0%) and caterpillars (63%) (Wilcoxon signed-rank test: control prey/caterpillars, p = 0.008). The GLM with the lowest AIC included bird behaviour (refused or eaten) as a response variable, bird age, sex, weight, and trial duration as fixed factors. No effect of bird age or sex on attack probability, but a strong effect of trial duration on refuse rate, were found, as expected, since the prey was considered refused after 20 minutes (p < 0.001, Table S1). A separate linear model showed a strong correlation of refuse rate and weight (refuse rate ∼ weight; p = 0.001), heavier birds tending to show a lower refuse rate.

**Fig. 4:**
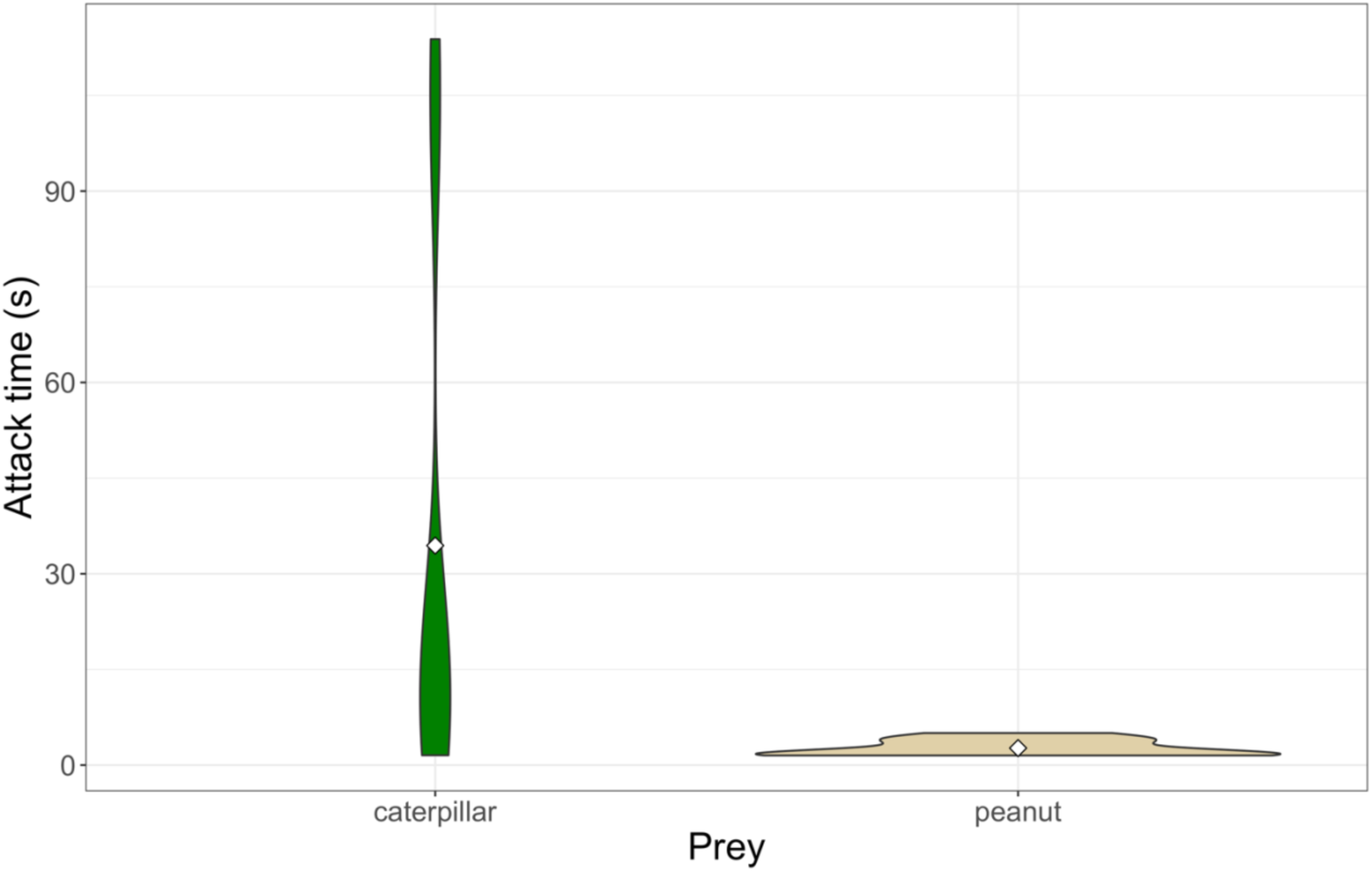
Violin plot showing the attack time in the palatability assay on caterpillars. *Attack time* differed markedly between control prey and caterpillars (Wilcoxon’s signed rank test, p < 0.001).

### Vision modelling

Chromatic contrast modelling suggests that the melanic morph cannot be differentiated from the camouflage net by birds, but it can be differentiated from the white morph. Both morphs can be differentiated from Box Tree Leaves (Fig. S1). The white morph is more conspicuous than the melanic morph both on camouflage net and on Box Tree leaves (Fig. S1). Achromatic contrast modelling suggests that both morphs can be differentiated on the camouflage net. On Box Tree leaves, however, the white is more conspicuous than the melanic on the upper leaf side, whereas on the underside the white morph is not distinguishable from Box Tree leaves (Fig. S2). The white morph has UV-reflective wings, but not the melanic morph. Reflectance spectra for Box Tree leaves, camouflage net and moth morphs can be found in Fig. S3.

### Predation

#### Preliminary analysis

Predation data were obtained using 1350 individual prey, tested with 45 birds. Moths were positioned randomly in the arena, and an absence of bias in moth positioning before the trials was confirmed (Chi-squared test, p = 0.935). In the linear model fitted with raw data, only cell C1 emerged as occupied slightly significantly more often than others (p = 0.045) (Table S2). However, cells in the centre of the arena received markedly more attacks than cells at the edge (Chi-squared test, p < 0.001, Table 1). The order of attack (i.e. whether the moth is attacked as first, second, etc.) didn’t explain attack probability for both morphs in the linear model (p = 0.86). We found, however, that searching time increases significantly with order (i.e., towards the end of the experience) (p < 0.001).

#### Comparison of different treatments

To see how *morph-specific attack rate* and random scenario are defined, please refer to Fig. 5 and to the Material and Methods section. In the treatment with equal morph ratios, *morph-specific attack rate* was 0.59 for the white and 0.41 for the melanic morph. Both are different from the 0.5 expected attack rate under random scenario (One Sample t-test, p = 0.002). Melanic moths experience a lower attack rate than white moths, and lower than the expected random rate (Fig. 5).

**Fig. 5:**
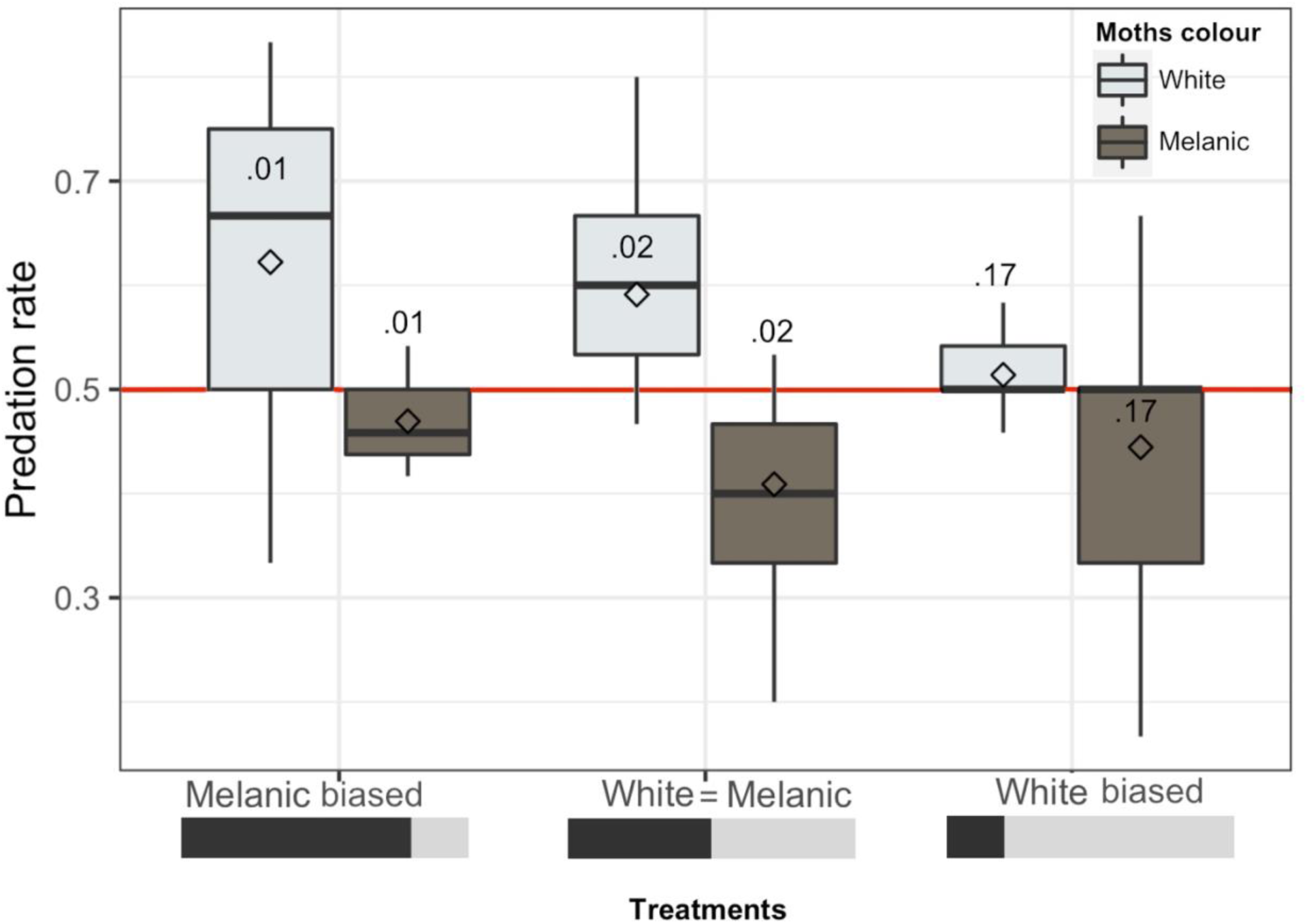
Box Plot showing the *attack rate* in the predation experiment. White squares represent the median, black bars the mean and the red line the morph-specific attack rate expected under a random scenario. P values from Wilcoxon’s Signed Rank test comparing observed and expected attack rates are shown below the bars. The *overall attack rate* was defined as the number of attacked moths of a certain morph over all moths presented, and the *morph-specific attack rate* as the number of attacked moths of a certain morph over available moths of that same morph. In a random scenario, birds would be expected to attack the moth morphs according to their relative abundance. For example, in the melanic-biased treatment (20:80 white melanic ratio), 6 out of 30 moths were whites and 24 melanic. In this treatment, considering that birds could eat 15 moths, the *overall attack rate* in a random scenario for white moths would correspond to the abundance of whites, 0.2, that corresponds to 3 out of 15 eaten moths (0.2*15 = 3). Since only 6 individuals were melanic, this corresponds to a *morph specific attack rate* of 0.5, that corresponds to 3 out of 6 available melanics (0.5*6 = 3). The same for the melanic morph: the *overall attack rate* would be 0.8 (corresponding to 0.8*15 = 12 moths) and the *morph-specific* one would be 0.5 (corresponding to 0.5*24 = 12 white moths).

In the melanic-biased treatment, the *morph-specific attack rate* was 0.62 for the white and 0.47 for the melanic morph. Both are different from the 0.5 attack rate expected in the absence of FDS and difference in detectability between morphs (random scenario; One Sample t-test, p = 0.002; see also Material and Methods), again with melanic moths experiencing a lower attack rate than white moths and compared to the random scenario (Fig. 5).

In the white-biased treatment, the *morph-specific attack* rate was 0.51 for the white and 0.44 for the melanic morph. Neither morph shows a significant deviation from the 0.5 expected rate under a random scenario (One Sample t-test, p = 0.17) (Fig. 5).

#### Frequency-dependence

In the case of positive FDS, a difference should be observed between treatments in the *morph-specific attack rate*. The white morph, when abundant, experiences a lower attack rate compared to the treatment with equal ratios (0.51 vs 0.59, p = 0.008). The melanic morph, when abundant, is attacked more heavily than in the treatment with equal ratios (0.41 vs 0.47, p = 0.034). This suggests that the white morph is under positive, whereas the melanic is under negative FDS. When at low frequency, the white morph is attacked more often than in the treatment with equal ratios (0.59 vs 0.62, p = 0.030), but the melanic morph, when at low frequency, is not attacked less often than in the treatment with equal ratios (0.41 vs 0.44, p = 0.444). However, when analysing the morph at low frequency, the statistical power is limited due to the rare morph being represented by only 6 prey items in the arena. This is reflected by the large variance in the predation rates on the rarer morph (Fig. 5).

#### Linear model

Morph color emerged as highly significant in the model, together with grid position. Both morph (melanic) and position (edge) had a negative estimate value in the model (Table 1), suggesting that melanic moths are attacked less heavily than white, and that prey positioned at the edge of the grid were receiving fewer attacks.

## DISCUSSION

Our results show how predation and frequency dependence contribute to the maintenance of polymorphism in the Box Tree Moth (*Cydalima perspectalis*), having two colour morphs that differ in their level of conspicuousness. Adults appear fully palatable to birds, whereas caterpillars appear to be defended and were generally avoided. Our results highlight that the melanic morph benefits from reduced predation compared to the white morph, and that it is best off when rare, whereas the white is best off when common. This pattern of opposite FDS acting on the two morphs brings new insight into the forces acting on prey phenotypes, and may contribute to the polymorphism observed in natural populations.

### Two life stages, two different strategies against predation

Adult Box tree Moths were highly palatable to blue tits, in agreement with the absence of alkaloids in their tissues (Leuthard et al. 2013), as well as with observed bird predation events (Brua, 2014) and photographs from citizen science repositories (unpublished data). Caterpillars, in contrast, were moderately unpalatable to blue tits in our experiments. Larvae feeding on box tree leaves may derive some chemical defence against predation, but this defence is not passed on to adults, suggesting that the two stages are exposed to different predation regimes. However, bird avoidance of caterpillars was highly variable. Bird age or sex may influence attack propensity on unpalatable prey (Gordon et al., 2021) but there was no effect of those factors in our analyses. Birds might also balance unpalatability with nutrition capacity (Barnett et al., 2007; Halpin et al., 2014). In our experiments, all birds were fed before the trial, and lighter birds showed an increased refuse rate, arguing against the idea that risk taking is fostered by poorer body condition. Nevertheless, caterpillars might be expected to represent attractive prey to blue tits in the middle of winter, perhaps explaining the tolerance of some birds to chemically-defended prey (Halpin et al., 2014).

### Polymorphism in the Box-Tree Moth

Prey polymorphisms attract attention in evolutionary ecology because they stand as excellent models to get insight into the functional role of character ecology in evolution (Alatalo and Mappes, 1996; Bond, 2007). Much attention was given to the operation of selection imposed by the visual and cognitive ecology of predators to understand, on the one hand, the maintenance of variation in the appearance of cryptic prey as a result of negative frequency dependence and, on the other hand, the evolution of conspicuousness in a context of aposematism, warning signals, or mimicry (Ruxton et al., 2004; Merilaita et al., 2017). Our study brings an interesting example of intermediary situation by evaluating the effect of predation on morphs that differ markedly in their conspicuousness but are overall highly palatable.

Our results suggest that the melanic morph of the Box Tree Moth may enjoy a survival advantage in the face of predation by insectivorous birds. This advantage likely owes to the melanic morph being less visible than the white morph. As expected for a morph relying on crypsis, this advantage is stronger when the melanic morph is rare. By contrast, the more conspicuous white morph does not appear to have a symmetrical advantage when rare, and is best off when common. The two morphs are therefore subject to different types of frequency-dependent predation in our experiments.

Palatable prey are expected to be under negative frequency-dependent predation because predators get better at detecting cryptic prey as they become more abundant (familiarity effect). However, these expectations may vary owing to the level of prey visibility, because predator familiarity is not expected to affect detection probability for conspicuous prey types (Bond and Kamil, 2002). Positive frequency-dependent predation is often observed in the context of aposematism and predator learning, but it is not usually reported for palatable prey. Here, the observed positive FDS in the conspicuous morph of this palatable moth may occur in the context of a dilution effect (Lindström et al., 1997; Cresswell and Quinn, 2011).

Dilution effect may be found for palatable, conspicuous prey living in aggregations, since they are easily detected both in small and large groups, and so no negative effect of gregariousness is expected (Bond and Kamil, 2002). However, in larger groups the per capita risk of attack decreases with increasing group size (Wrona, 1991; Lehtonen & Jaatinen, 2016). Here, although the moths were not presented in groups, they were found repeatedly by the birds over a relatively short period of time, which could reduce individual predation risk as the density in the arena increases. Our data shows that positive FDS can arise for conspicuous and palatable prey irrespective of group living.

### Are birds driving frequencies of the box tree moth in nature?

Bird predation is an important source of selection on butterflies and moths and may influence morph frequencies in a balance with other processes (Bowers et al., 1985; Nokelainen et al., 2012; Cook and Saccheri, 2013; Chouteau et al., 2017). Across Europe, morph frequencies in the Box Tree Moth are relatively stable, with the melanic morphs being found at a frequency of 15-25% (unpublished data), suggesting that some process may stabilise frequencies. A balance between crypsis in the melanic and dilution in the white morphs may contribute to the maintenance of the morphs, limiting the frequency of the black morph but also preventing its loss. However, it is unclear how a stable equilibrium may be derived from the interplay of those processes, so how those forces oppose or not at intermediate frequencies should be investigated further. Moreover, the design with a fixed number of prey attacks may lead to the opposite frequency-dependent effects partly to hide each-other. Trials limited by time and not by the number of prey attacked may allow to explore this aspect.

As in other prey displaying colour polymorphisms (e.g., Lepidoptera (Nokelainen et al., 2012; Chouteau et al., 2017), *Harmonia axyridis* (Michie et al., 2010), *Podarcis* lizards (Pérez i de Lanuza et al., 2017), birds (Galeotti et al., 2003), etc.) visual predation may be only one of the drivers maintaining polymorphism. Differences in mating success between the two morphs, in chemical noxiousness in their larvae, or linked deleterious effects near the loci controlling them, are candidate processes which may contribute to the polymorphism (Wang, 2011; Gordon et al., 2015; Chouteau et al., 2016; Jay et al., 2019).

Invasive species may reach spectacular densities. During population outbreaks, the Box tree moth reaches densities at which predators are unlikely to influence morph frequencies (Ledru et al., 2022; Poloni, unpublished data). However, once caterpillars have defoliated all available hosts, moths populations may persist at lower densities according to the availability of foodplants (Suppo et al., 2020). During such periods selection on morph frequencies may be operated by visual predators.

Other factors may of course also influence attack rates in our experiments, as they do in other species, like behavioural escape strategies (Svensson et al. 2003) or wing UV reflection and transparency, that increase and decrease detectability for predators, respectively (Henze et al., 2018; Chan et al., 2019). The wings of the Box Tree Moth are both UV reflective (white morph) and diaphanous, with possible effect on crypsis efficiency. Follow-up experiments could therefore use natural daylight, and perhaps more natural substrate structure may allow to explore these factors further.

As in any specific study system, all these mechanisms may be intertwined, together with invasion processes, in how they influence wing colour polymorphism. Our study shows, with a natural system, that morphs with drastic different levels of conspicuousness may be influenced by predation in opposite directions. In particular, a palatable conspicuous morph appears to show positive FDS, whereas its less conspicuous counterpart follows negative FDS. A better understanding of the genetic and ecological factors associated with this polymorphism is necessary to better understand how a balance of mechanisms within and between populations produces equilibrium frequencies and fosters the maintenance of diversity in an invasive insect.

## AUTHORS CONTRIBUTION

RP and MJ conceived the study; RP, JM, MJ and ON, designed the experiments; RP, MD and ON collected data for predation experiments; RP and MD analysed the data; RP and MJ wrote the manuscript. All authors contributed to the discussion of the results and provided their contribution to the manuscript.

## ACKNOWLEDGEMENTS

We are deeply indebted with all people that helped us during experiments in the Konnevesi Research Station, in particular to Helinä Nisu for her expertise on bird handling and training and to Sanni Silvasti, for being the best Cicero we could have in station. We are also thankful to Doris Gomez who gave us useful advice about spectrometry data, to Céline Teplitsky and Gaël Thébaud foradvice on linear models, and to Yann Le Poul and Pierre Lacoste for help with statistical analysis. This project was funded by ANR-18-CE02-0019-01 and ERC-StG-243179 grants to MJ, and the Academy of Finland, project no. 345091 to JM.

## DATA AVAILABILITY

Raw data and scripts used for this paper can be found in supplementary files and at https://github.com/xxx (all data and scripts will be shared upon publication).

## CONFLICT OF INTEREST

The authors declare no conflicts of interest.

## SUPPLEMENTARY MATERIAL

Supplementary methods: more detailed description of methods used for this paper. Supplementary material: supplementary tables and figures.

Supplementary data: raw data and scripts used for analysis.

## Notes

### Competing Interest Statement

The authors have declared no competing interest.

