## Supplementary Methods for "Positive and negative frequency-dependent selection acting on polymorphism in a palatable moth"

### Model species

Box tree moth specimens were obtained from a laboratory stock founded in 2020 at the Centre d'Ecologie Fonctionelle et Evolutive (France) using wild populations from Saint-Clément-de-Rivière, France (GPS coordinates: 3.84, 43.71) and Le Caylar, France (GPS coordinates: 43.86, 3.32). Caterpillars were fed with fresh branches of *Buxus sempervirens* collected from the same localities. Adults and caterpillars were frozen alive at -80 °C and shipped on dry ice to the Konnevesi Research Station (Central Finland).

Wild blue tits (*Cyanistes caeruleus*) were caught from baited traps at Konnevesi Research Station, where all experiments took place, in February-March 2022. Once trapped, all birds were measured, weighted, aged, sexed, and housed individually in plywood cages (80 cm x 65 cm x 50 cm) with a daily light period of 11L∶13D as described by Ham et al. (2006). Birds were fed sunflower seeds, peanuts, and vitamin-enriched tallow and provided with fresh water ad libitum. Cages were regularly cleaned. After the experiment, all birds were ringed and released at the capture site..

### Visual modeling

Spectrometer measurements and visual modelling were used to model how birds perceive the background chosen and the two morphs of the Box Tree Moth. The measurements were taken using a Maya 2000-PRO spectrometer model MAYP11351 (Ocean Optics, Dunedin, FL), with automatic averaging of three scans, automatic integration times, and by enabling the "electric dark correction" option.

Reflectance measures were taken between 300 and 700 nm. 5 white and 5 melanic moths were scanned, 1 measure per wing side in the melanic individuals and 2 per side for white individuals, corresponding to the white and melanic part of the wings. The camouflage net used as a background was also measured, doing 5 recordings on different spots both for green and brown patches. Finally, *Buxus sempervirens* leaves were measured, recording 15 leaves with one measure for each side of the leaf. Then, chromatic and achromatic contrasts were calculated using the package pavo 2 (Maia et al., 2019) in units of Just Noticeable Differences (JND) between moths and the camouflage net, as well as between moths and box tree leaves. To do so, spectra were first smoothed to remove background noise. To predict if the colours are distinguishable by the receiver, we used then the function "vismodel" using the discriminability model of Vorobyev & Osorio (1998). Cone density values were based on data available for the blue tit, *Cyanistes caeruleus*, namely: 4 cones with density equal to 1:1.92:2.68:2.7 (UV: small:medium:long wavelength) (Hart et al., 2000)) and Weber fraction, chromatic = 0.1 (Meyer & Bowmaker, 1993), achromatic = 0.2 (Lind et al., 2013, Silvasti et al., 2021). Chromatic and achromatic distances were finally obtained using the function "bootcoldist" with the above parameters.

**Predation experiments**

The day before trial birds were provided with 4 moths (2 of each morph) in their home cages, to familiarize with the prey. Then, birds were habituated to the aviary by letting them overnight in groups of 5 and forage palatable, familiar food from the floor (peanuts and sunflower seeds). Trials were run during the same day or the following day. Prior to the start of each trial, the experimental bird was food-deprived for 1-1:30 h to ensure that it was motivated to forage.

Each trial was observed through a one-way mirror and the timing, order, and position of each attack were recorded, as well as other bird behaviours, such as beak wiping or cleaning. The experiment was terminated when the bird had eaten half (15) moths.
