## Supplementary Material for "Positive and negative frequency-dependent selection acting on polymorphism in a palatable moth"

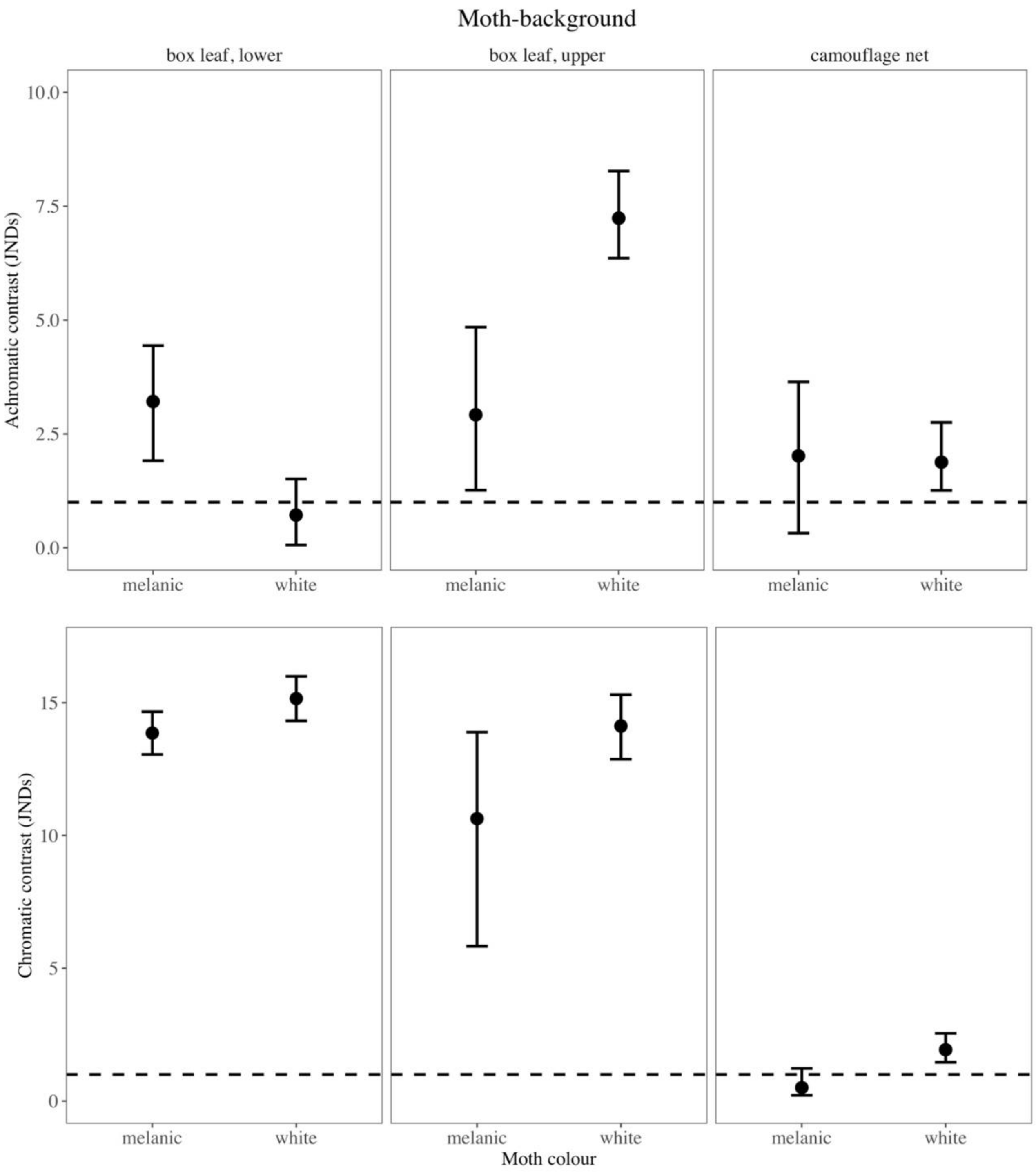


Fig. S1: Chromatic and achromatic contrasts measured in Just Noticeable Difference (JND) between the two morphs of the Box Tree Moth and Buxus leaves or between moths and the camouflage net used in the experiment.

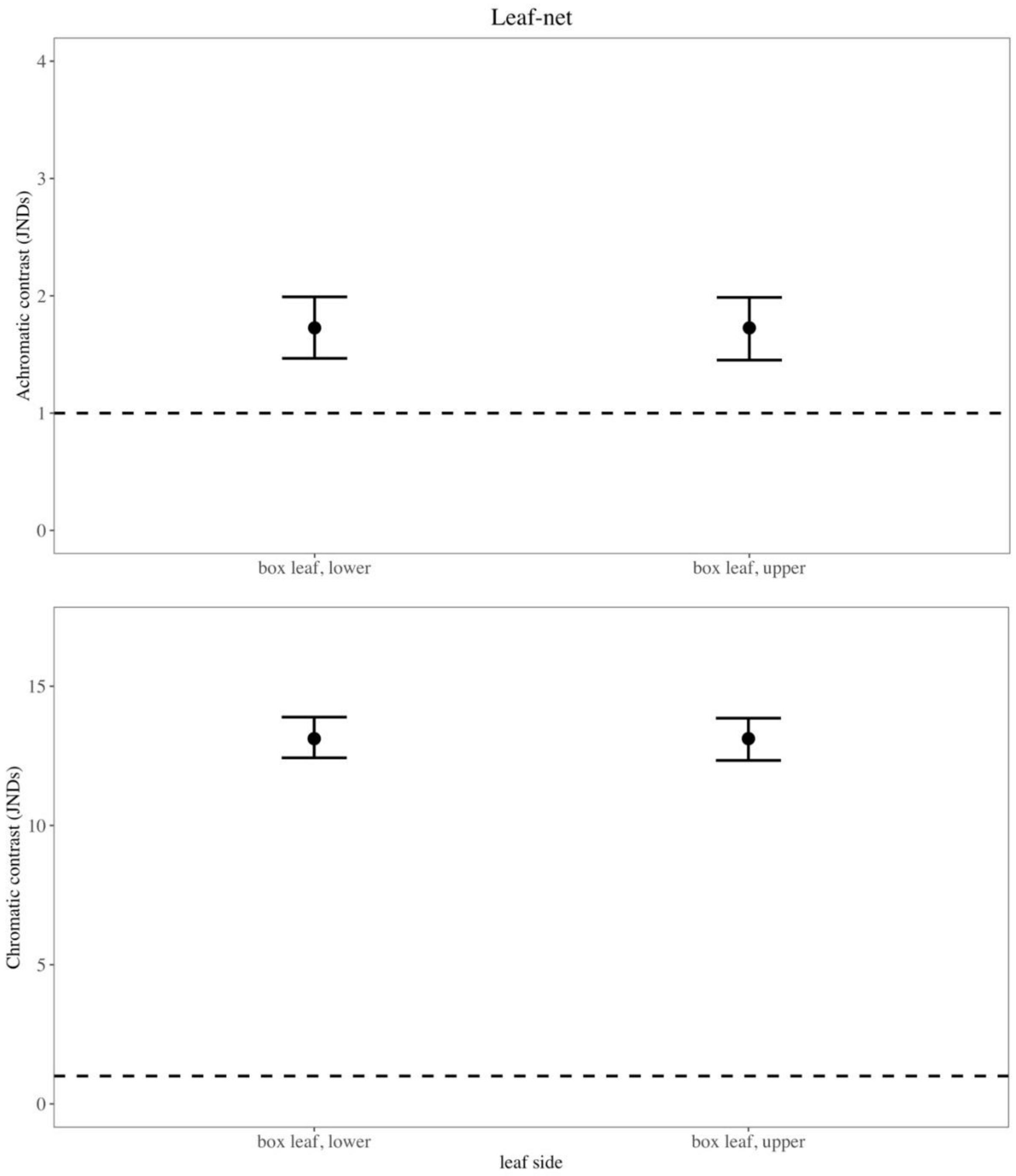


Fig. S2: Chromatic and achromatic contrasts measured in Just Noticeable Difference (JND) between Buxus leaves and the camouflage net used in the experiment.

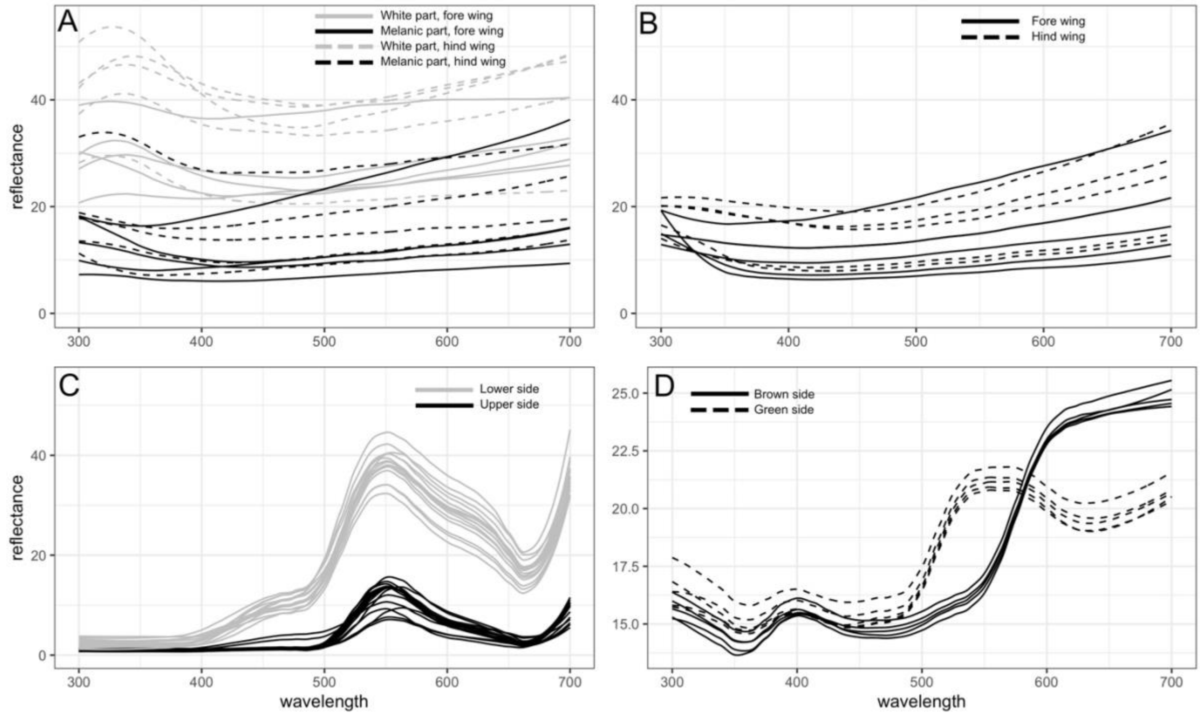


Fig. S3: Reflectance spectra for white Box Tree Moth (A), melanic Box Tree Moths (B), *Buxus sempervirens* leaves (C) and camouflage net (D).

| | E | SE | Z | P |
|---|---|---|---|---|
| **(Intercept)** | 2,144 | 4,461 | 0,481 | 0,631 |
| **age (juv)** | -0,627 | 0,503 | -1,247 | 0,212 |
| **Sex (male)** | 1,224 | 0,568 | 2,155 | 0,031 |
| **Weight_In** | -0,555 | 0,383 | -1,449 | 0,147 |
| **Trial Duration** | 0,063 | 0,011 | 5,629 | 0,000 |

Table S1: Fixed effects for the generalized linear model performed with data on refuse rate for caterpillars, including morph, treatment and grid position (edge or center) as fixed factors (AIC = 1835).

| | E | SE | Z | P |
|---|---|---|---|---|
| (Intercept) | 1,253 | 0,359 | 3,494 | 0,000 |
| Grid Position=A2 | -0,241 | 0,492 | -0,490 | 0,624 |
| Grid Position=A3 | 0,134 | 0,517 | 0,258 | 0,796 |
| Grid Position=A4 | 0,000 | 0,507 | 0,000 | 1,000 |
| Grid Position=A5 | -0,458 | 0,482 | -0,950 | 0,342 |
| Grid Position=A6 | 0,000 | 0,507 | 0,000 | 1,000 |
| Grid Position=A7 | -0,352 | 0,487 | -0,723 | 0,469 |
| Grid Position=A8 | -0,124 | 0,499 | -0,249 | 0,803 |
| Grid Position=B1 | 0,279 | 0,530 | 0,526 | 0,599 |
| Grid Position=B2 | 0,000 | 0,507 | 0,000 | 1,000 |
| Grid Position=B3 | 0,439 | 0,546 | 0,804 | 0,421 |
| Grid Position=B4 | -0,241 | 0,492 | -0,490 | 0,624 |
| Grid Position=B5 | -0,124 | 0,499 | -0,249 | 0,803 |
| Grid Position=B6 | -0,458 | 0,482 | -0,950 | 0,342 |
| Grid Position=B7 | -0,241 | 0,492 | -0,490 | 0,624 |
| Grid Position=B8 | -0,241 | 0,492 | -0,490 | 0,624 |
| Grid Position=C1 | -0,939 | 0,469 | -2,004 | 0,045 |
| Grid Position=C2 | 0,619 | 0,566 | 1,093 | 0,274 |
| Grid Position=C3 | -0,124 | 0,499 | -0,249 | 0,803 |
| Grid Position=C4 | -0,560 | 0,478 | -1,171 | 0,242 |
| Grid Position=C5 | -0,124 | 0,499 | -0,249 | 0,803 |
| Grid Position=C6 | 0,279 | 0,530 | 0,526 | 0,599 |
| Grid Position=C7 | -0,241 | 0,492 | -0,490 | 0,624 |
| Grid Position=C8 | 0,279 | 0,530 | 0,526 | 0,599 |
| Grid Position=D1 | 0,134 | 0,517 | 0,258 | 0,796 |
| Grid Position=D2 | -0,458 | 0,482 | -0,950 | 0,342 |
| Grid Position=D3 | -0,241 | 0,492 | -0,490 | 0,624 |
| Grid Position=D4 | -0,352 | 0,487 | -0,723 | 0,469 |
| Grid Position=D5 | 0,134 | 0,517 | 0,258 | 0,796 |
| Grid Position=D6 | -0,352 | 0,487 | -0,723 | 0,469 |
| Grid Position=D7 | -0,124 | 0,499 | -0,249 | 0,803 |
| Grid Position=D8 | -0,352 | 0,487 | -0,723 | 0,469 |
| Grid Position=E1 | 0,439 | 0,546 | 0,804 | 0,421 |
| Grid Position=E2 | -0,560 | 0,478 | -1,171 | 0,242 |
| Grid Position=E3 | -0,241 | 0,492 | -0,490 | 0,624 |
| Grid Position=E4 | 0,134 | 0,517 | 0,258 | 0,796 |
| Grid Position=E5 | 0,000 | 0,507 | 0,000 | 1,000 |
| Grid Position=E6 | 0,134 | 0,517 | 0,258 | 0,796 |
| Grid Position=E7 | -0,458 | 0,482 | -0,950 | 0,342 |
| Grid Position=E8 | -0,560 | 0,478 | -1,171 | 0,242 |

Table S2: Fixed effects for the generalized linear model performed with predation data, testing for a possible position bias.
